# Hippocampal reactive neural stem cells are able to phagocytose and have an immunological molecular signature

**DOI:** 10.1101/2024.01.26.577346

**Authors:** Oihane Abiega, Rodrigo Senovilla-Ganzo, Maroua El-Ouraoui, Teresa Muro-García, Lorena Ruiz-Clavijo, Fernando García-Moreno, Carlos Fitzsimons, Soraya Martín-Suárez, Juan M. Encinas

## Abstract

Hippocampal neural stem cells (NSCs) are the drivers of neurogenesis in the dentate gyrus (DG) of most mammals including humans. During neuronal hyperactivity NSCs become reactive NSCs (react-NSCs), characterized by their activation, morphological changes, and symmetric division, abandoning their neurogenic programme and transforming into reactive astrocytes. Here, using different pathological models that induce react-NSCs in the DG, we looked for novel features of react-NSCs both histologically and by total RNA sequencing. We report that in two pathological models were react-NSCs emerge (mesial temporal lobe epilepsy (MTLE) and traumatic brain injury (TBI)) react-NSCs are capable of phagocytosis of dead cells, a typical immunological function carried out mainly by microglia in the brain. Importantly, MTLE-induced react-NSCs show phagocytic function in tissue and a predominantly immunological molecular signature, with a broad upregulation of phagocytosis-related gene expression. Our results describe a new function of react-NSCs as phagocytic and immunologically active cells in the hippocampal neurogenic niche.

## INTRODUCTION

Hippocampal neurogenesis or the process of creation of new neurons in the subgranular zone (SGZ) of the hippocampal dentate gyrus (DG), is a key process involved in memory formation, pattern separation, learning, and mood disorders like fear, anxiety and stress (Clelland et al. 2009; Deng et al. 2009; Dupret et al. 2008; Farioli-Vecchioli et al. 2008; Saxe et al. 2006). Hippocampal neural stem cells (NSCs) are the drivers of neurogenesis. In physiological conditions a small proportion of NSCs get activated and divide asymmetrically, giving rise to amplifying neural progenitors (ANPs) that will eventually become new mature neurons that will get integrated into the hippocampal circuitry.

After a series of symmetric divisions, NSCs differentiate into astrocytes (Sierra et al. 2015). However, in conditions of neuronal hyperactivity, such as in a model of mesial temporal lobe epilepsy (MTLE) induced by intracerebral injection of Kainic acid (KA), the NSC pool gets massively activated and NSCs become reactive NSCs (react-NSCs), characterized by switching to a symmetric division to create more react-NSCs instead of ANPs, thus abolishing neurogenesis (Sierra et al. 2015). Moreover, react-NSCs differentiate into reactive astrocytes, a cellular type linked to astrogliosis which is one of the hallmarks of MTLE (Sierra et al. 2015). Importantly, this conversion to react-NSCs happens in two different models of KA administration (intra-hippocampal and intra-amygdalar), demonstrating that react-NSCs arise due to neuronal hyperactivity and not to a local effect of KA (Sierra et al. 2015; Muro-García et al. 2019). Importantly, react-NSCs have also been found to arise in other pathological conditions such as traumatic brain injury (TBI), thus proving that these type of cells are not specific of MTLE models.

Phagocytosis, or the process of engulfing different types of cellular debris, is fundamentally an immunological process that is predominantly carried out by microglia in the brain. In the hippocampal neurogenic niche, apoptosis (programmed cell death) of ANPs has been shown to be coupled to microglial phagocytosis (Sierra et al. 2010), making it a key process for the maintenance of the niche. Interestingly, other cells such as astrocytes and neuroblasts are also capable of performing phagocytosis in the brain (Abiega et al. 2016), albeit at much lower numbers and efficiency than microglia.

In the present study we asked whether hippocampal react-NSCs could have extra-neurogenic roles in the maintenance of the neurogenic niche. We also asked whether react-NSCs could have differences in their transcriptome that could be linked to possible extra-neurogenic functions. Using LPA_1_-EGFP mice (where NSCs and react-NSCs are specifically labelled in green) for total RNA sequencing and LPA_1_ and nestin-GFP mice for in tissue assessment of apoptotic cell phagocytosis, we studied the phagocytic capacity and molecular signature of NSCs and react-NSCs in a model of MTLE and TBI. We find that react-NSCs are capable of phagocytosing apoptotic cells in both pathological conditions, in contrast to NSCs which do not perform phagocytosis. We also show that this capacity for phagocytosis in react-NSCs is coupled to a predominantly immunological molecular signature, and to an up-regulation of various phagocytosis-related genes and gene pathways. Thus, we here report that phagocytosis is a new extra-neurogenic function of react-NSCs that contributes to the maintenance of the homeostasis of the hippocampal neurogenic niche.

## EXPERIMENTAL PROCEDURES

### Animals

All the experiments were carried out using Lysophosphatidic acid receptor 1–GFP (LPA1-GFP) (Gong et al. 2003) transgenic mice and nestin-GFP transgenic mice (Mignone et al. 2004). Both mouse lines were crossbred with C57BL/6 mice. All procedures were approved by the University of the Basque Country (UPV/EHU) Ethics Committees (Leioa, Spain) and Diputación Foral de Bizkaia under protocol M20/2015/236. All procedures followed the European directive 2010/63/UE and National Institutes of Health guidelines.

### Intrahippocampal injection of KA

2 months old (mo) mice were anesthetized using isoflurane and received a single subcutaneous shot of the analgesic buprenorphine (1mg/kg) (Buprecare, Animalcare Lted). Mice were positioned in the stereotaxic apparatus and a 0.6mm hole was drilled at the following coordinates taken from Bregma: anteroposterior (AP) −1.8mm, laterolateral (LL) −1.6mm. A glass microcapillary was then inserted at −1.9mm dorsoventral (DV) and 50nl of either phosphate buffered saline (PBS) or Kainic acid (KA, 20nM) were injected into the right hippocampus in 11 pulses (4,6nl each) using a microinjector (Nanoject II, Drummond Scientific, Broomal, PA, USA). After 2 minutes, the capillary was slowly retracted to avoid reflux and the mice were sutured. Then mice were left to awaken and recover at RT. Mice were closely monitored during the hours following surgery.

### Controlled Cortical Impact model of Traumatic Brain Injury

A moderate model of controlled cortical impact was used. In brief, mice were anesthetized using isoflorane and placed in a stereotaxic frame. Then the skull was exposed and a 4 mm x 4 mm square shaped craniotomy was opened using a drill, avoiding bleeding to ensure undamaged meninges. Then the impact piston (3mm diameter) of a Leica impactor was placed on the surface of the brain at an angle of −1 in the vertical axis and the brain was impacted using the following settings: 5,5 m/s velocity, 600 ms dwell time and 1mm depth. Following the impact, the removed skull was glued back to close the craniotomy, and the skin over the skull sutured. Finally, the animal was placed on a thermal blanket for recovery until fully awake. Sham mice underwent the same procedure without the impact.

### FACS Sorting

Neural Stem Cells (NSCs) were isolated from the injected hippocampi as described previously (Abiega et al. 2016). At 3 days post-surgery, 4 mice per experimental replica (total of 5 replicas per treatment) were sacrificed by cervical dislocation, brains isolated and the right (injected) dentate gyri (DG) dissected following previous protocols (Walker y Kempermann 2014). DGs were then placed in enzymatic solution (116 mM NaCl, 5.4 mM KCl, 26 mM NaHCO3, 1 mM NaH2PO4, 1.5 mM CaCl2, 1mM MgSO4, 0.5 mM EDTA, 25 mM glucose, 400mM L-cysteine) containing papain (20 U/ml) and DNAse I (150 U/μl, Invitrogen), minced and placed for 10 minutes at 37°C to be digested. Then samples were mechanically homogenized using a pipette and filtered through a 40 μm nylon strainer to a Falcon tube containing 5 ml of 20% heat inactivated FBS in HBSS. To further clean the samples, myelin was removed using Percoll gradients. For this, cells were centrifuged (200 G, 5 min) and resuspended in a 20% Solution of Isotonic Percoll (20% SIP; in HBSS), obtained from a previous stock of SIP (1/10 PBS 10X in Percoll). Then, each sample was very gently layered with HBSS (to avoid mixing with SIP) and centrifuged (200 G, 20 min) with minimum acceleration and no brake to avoid disruption of the interphase. Then the interphase was removed, cells washed in HBSS by centrifugation (200 G, 5 min) and the pellet was resuspended in 500 μl of sorting buffer (25 mM HEPES, 5 mM EDTA, 1% BSA, in HBSS). NSC sorting was performed by FACS Jazz (BD). Samples were stained for 10 min with propidium iodide (PI) to avoid sorting dead or dying cells. Each experimental sample was ran alongside its respective negative controls in order to establish proper gatings: WT DG homogenate as a control for GFP fluorescence, and experimental sample without PI as a control for dead cells. Samples were gated to exclude debris, doublets, dead cells, and non-GFP+ cells. GFP+ cells were sorted in RLT buffer containing β-mercaptoethanol, were vortexed immediately after sorting and stored at −80°C until further processing.

### RNA isolation

Total RNA was thawed on ice and isolated using a Norgen single cell RNA purification kit (Cat. 51800, Norgen Biotek, Ontario, Canada) following manufacturer’s instructions, including a DNAse treatment step. Purified RNA was quantified in a Nanodrop 2000.

### Library preparation and sequencing

To perform RNA-seq samples were sent to the Genomic Platform at CICbioGUNE (Derio, Spain). The RNA quantity and quality was assessed using Qubit RNA assay kit (Invitrogen) and Agilent RNA 6000 Pico Chips (Agilent Technologies, Cat.# 5067-1513). All samples presented enough concentration and integrity to perform the experiments. Sequencing libraries were prepared using “SMARTer Stranded Total RNA-seq Kit v3 – Pico Input Mammalian” kit (Takara Bio, Cat. # 634487) and the “SMARTer RNA Unique Dual Index Kit - 24U” (Takara Bio, Cat. No. 634451), following “SMARTer Stranded Total RNA-seq Kit v3 – Pico Input Mammalian User Manual”. The protocol was started with 1.5 - 2 ng of good quality total RNA (RNA concentration was estimated by Bioanalyzer). Briefly, when needed RNA was fragmented and 1st strand cDNA synthesis was performed using SMARTScribe reverse Transcriptase, for 180 min at 42°C, 10 min at 70°C and pause at 4°C. Afterwards, Illumina Adapters containing Unique Dual Indexes were added, in a preamplification PCR (60 sec at 94°C; 5 cycles of 15 sec at 98°C, 15 sec at 55°C, 30 sec at 68°C; and pause at 4°C). Then, Ribosomal cDNA was depleted with ZapR v3 and R-Probes v3. Finally, enrichment of libraries was achieved by PCR (60 sec at 94°C; 14 * cycles of 15 sec at 98°C, 15 sec at 55°C, 30 sec at 68°C; and pause at 4°C). Afterwards, libraries were visualized on an Agilent 2100 Bioanalyzer using Agilent High Sensitivity DNA kit (Agilent Technologies, Cat. # 5067-4626) and quantified using Qubit dsDNA HS DNA Kit (Thermo Fisher Scientific, Cat. # Q32854). Raw images produced by the Illumina sequencer are generated using sequencing control software for system control and base calling, through the integrated primary analysis software known as RTA (Real Time Analysis). The binary output containing base calls (BCL) is subsequently transformed into FASTQ files using Illumina Inc.’s bcl2fastq package (FastQC, Andrews et al 2010).

### Bioinformatic analysis

The quality of sequenced FASTQs was evaluated using FASTQC (Andrews et al 2010) and the first 14 nt of Read2 were trimmed as indicated by TakaraBio with *cutadapt* (Marcel Martin et al, 2011). The alignment was carried out with STAR v2.7.3a (Dobin et al. 2013) against Ensembl genome of Mus musculus (dna.primary_assembly ang gtf v109). To obtain the count matrix, htseq-count (v 1.99.2, -s reverse) (Anders, Pyl, y Huber 2015) was run with strand specificity and imported to R v4.2.2, where it was analysed with DESeq2 (Love, Huber, y Anders 2014). DESeq2 provides methods for differential expression tests based on negative binomial generalized linear models. There was only one experimental factor: ∼ Treatment (KA vs Control). The differentially expressed genes were selected based in the p.adjusted (Benjamini-Hochberg) value < 0.05. For data visualization and functional enrichment: ggplot2 (Wickham 2016) and clusterProfiler (Wu et al. 2021; Yu et al. 2012) were used. The deconvolution analysis was carried out based on the single cell transcriptomic data from the published dataset from (Hochgerner et al. 2018); and the previous exploratory analysis was carried out with Seurat (Stuart et al. 2019) in R. The selected dataset contained RGL and astrocytes extracted from the global dataset (all unfiltered cell types) based on their annotation (RGL, Astro-adult) of the more advanced stages available (P120, P132). Then, the dataset was normalized with default *Seurat* SCTransform parameters. This SCT normalised assay was used to deconvolute our RNA-seq data with deconvolute tool from granulator package (R) and the method svr (Newman et al. 2015). The code used can be found in https://github.com/rodrisenovilla/EncinasLab. Analysed data will be uploaded conveniently at GEO repository (https://www.ncbi.nlm.nih.gov/geo/) once they are published.

### Immunofluorescence

Experimental procedures followed previously established protocols with minor modifications optimized for transgenic mice (Encinas et al. 2011). Briefly, animals underwent transcardial perfusion with 25 ml of PBS, succeeded by 25 ml of 4% (w/v) paraformaldehyde (PFA) in PBS (pH 7.4). Brains were extracted and postfixed in 4% PFA for 3 hours at room temperature (RT), followed by transfer to PBS and storage at 4°C. Serial sagittal sections, each 50 µm thick, were obtained using a Leica VT 1200S vibrating blade microtome (Leica Microsystems GmbH, Wetzlar, Germany). Slices were systematically collected via random sampling, with the right hemisphere sagittally sliced in a lateral-to-medial direction, including the entire DG. The 50 µm slices were collected in 6 parallel sets, each set comprising 12 slices, with 300 µm of distance between subsequent slices. One of these series was immunostained and used for imaging and quantification purposes. Immunostaining procedures adhered to a standard protocol: sections were treated with a blocking and permeabilization solution (PBS containing 0.25% Triton-100X and 3% BSA) for 3 hours at room temperature. Following this, sections were incubated overnight at 4°C with primary antibodies diluted in the same solution. Post-incubation, sections underwent thorough washing with permeabilization solution (PBS containing 0.25% Triton-100X), followed by a 3-hour incubation at RT with fluorochrome-conjugated secondary antibodies diluted in the blocking solution. Following another round of washing, sections were mounted on gelatin-coated slides with DakoCytomation Fluorescent Mounting Medium (DakoCytomation, Carpinteria, CA). To enhance and facilitate the visualization of the GFP signal from transgenic mice, an antibody against GFP was employed. The antibodes used included chicken anti-GFP (Aves Laboratories, Tigard, OR) at a 1:750 dilution; goat anti-GFAP (DakoCytomation) at 1:1000; rabbit anti-S100β (DakoCytomation) at 1:750; AlexaFluor 488 donkey anti-chicken (Molecular Probes, Eugene, OR) at 1:500; AlexaFluor 568 donkey anti-goat (Molecular Probes) at 1:500; AlexaFluor 647 goat anti-rabbit (Molecular Probes) at 1:500. DAPI at 1:1000 (Sigma) served to stained cell nuclei.

### Image and video capture

Fluorescence immunostaining images were acquired using a Leica SP8 (Leica, Wetzlar, Germany) confocal microscope, using the manufacturer’s software. Subsequently, standard adjustments to brightness, contrast, and background were uniformly applied to the entire image dataset, without further modification using the Leica LAS X software. Shown images are z-stack projections. Orthogonal views of phagocytic pouches and 3D videos were made from z-stacks.

### Cell and phagocytosis quantification

For the quantitative analysis of apoptotic cell death density and % of phagocytic NSCs the following criteria were applied. Apoptotic cells were defined were based on previous criteria (Abiega et al. 2016), using nuclear morphology after DAPI staining, and identifying cells in which the chromatin structure (euchromatin and heterochromatin) was lost and appeared condensed and/or fragmented (pyknosis/karyorrhexis). NSCs were identified adhering to previously established criteria (Encinas et al. 2011), characterized as Nestin-GFP, GFAP-positive cells in LPA_1_-GFP mice, and as Nestin-GFP, GFAP-positive and S100β negative cells in Nestin-GFP mice. They were additionally defined as radial glia-like cells, situated in the subgranular zone (SGZ) or the lower third of the granule cell layer (GCL), with processes extending from the SGZ towards the molecular layer. Phagocytosis performed by NSCs was characterized following previous criteria (Abiega et al. 2016) as the formation of an enclosed, three-dimensional pouch of NSC processes surrounding an apoptotic cell.

Quantitative assessments of apoptosis, NSCs, and phagocytosis were conducted using unbiased stereology methods, as previously outlined, (Sierra et al. 2010) using Leica LAS X software. At least one immunostained series was imaged using a 40x oil immersion objective and 0,7µm step-sized z-stacks were made from the whole GCL of each slice. Total numbers of apoptotic cells, NSCs, and phagocytic NSCs were quantified for each z-stack, excluding those in the uppermost focal plane and in the upper and left edges of the stack. The values were normalized to the quantified volume of the SGZ+GCL for each animal. The total volume was obtained by measuring the area and thickness of the SGZ+GCL in each z-stack and the density of apoptotic cells and NSCs per mm^3^ was obtained.

### Statistical analysis

GraphPad Prism was used for statistical analysis. For the analysis of apoptotic cell density, Shapiro-Wilk test was used to check for normality and then significance was analysed using an unpaired t-test. For the analysis of % of phagocytic NSCs normality was checked using a Shapiro-Wilk test and significance was analysed using a one sample t-test. Only p<0.05 is reported to be significant. Data are shown as mean ± SEM (standard error of the mean).

## RESULTS

### Hippocampal NSCs and react-NSCs phagocytose apoptotic cells in MTLE in LPA_1_-GFP mice

First, we wanted to analyze phagocytosis of NSCs in tissue in a model of MTLE known to induce the change of “physiological” NSCs to react-NSCs. To induce MTLE, we injected Kainic acid (KA) in the hippocampus of 2mo mice (Sierra et al. 2015). This model reproduces the main hallmarks of human MTLE: seizures and hippocampal sclerosis (cell death, granule cell layer (GCL) dispersion, and inflammation) (Babb et al. 1995; Bouilleret et al. 1999; Heinrich et al. 2006; Kralic, Ledergerber, y Fritschy 2005; Nitta et al. 2008). To specifically visualize NSCs and react-NSCs, we used LPA_1_-GFP transgenic mice, where GFP is specifically expressed in NSCs (Walker et al. 2016). To test whether the KA injection was successful, we visually confirmed the induction of react-NSCs in the GCL and SGZ by immunofluorescence at 3dpKA (Sierra et al. 2015). Then we proceeded to quantify the phagocytosis of apoptotic cells by NSCs and react-NSCs (pyknotic nuclei stained with DAPI and surrounded by GFP and GFAP) (Abiega et al. 2016) in the GCL and SGZ of the DG (Figure 1A). Our quantifications show a significant increase in apoptotic cell numbers (Figure 1B) and 0,10% of phagocytic react-NCSs at 3dpKA as compared to no phagocytic NSCs in C mice (Figure 1C). Thus, only react-NSCs phagocytose apoptotic cells at 3dpKA.

**Figure 1.**
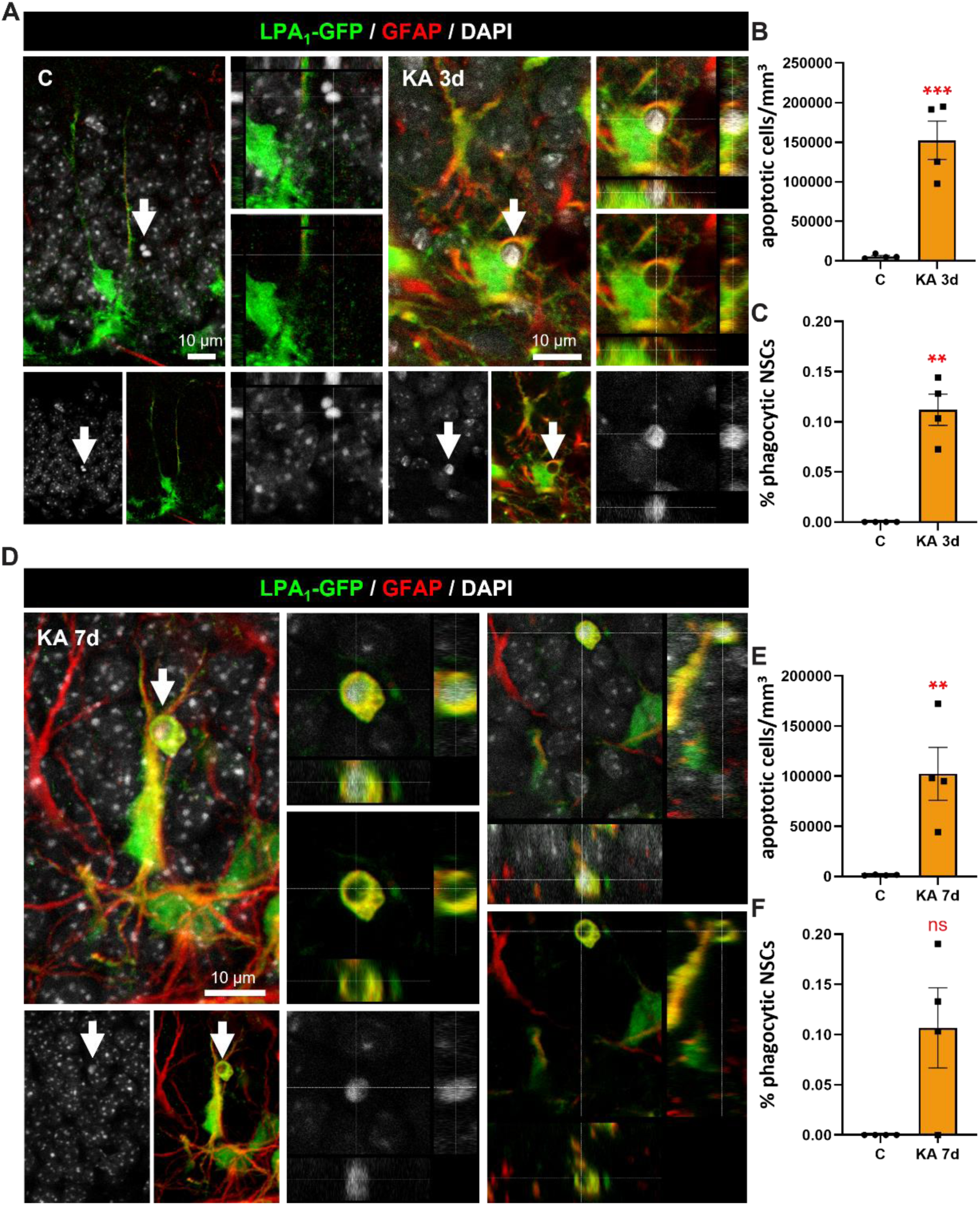
Hippocampal react-NSCs phagocytose apoptotic cells in a model of MTLE in LPA1-GFP mice. (A) Confocal microscopy images (projection from z-stacks) showing a non-phagocytic NSC (GFP+, GFAP+) and an apoptotic cell (pyknotic, stained with DAPI) in control and a phagocytic react-NSC (GFP+, GFAP+) engulfing an apoptotic cell (KA 3d). A high magnification orthogonal view is shown in the right panel for each treatment. (B) Quantification of apoptotic cells per mm^3^ in the GCL and SGZ of the DG at 3dpKA. (C) Quantification of the % of phagocytic NSCs at 3dpKA. (D) Confocal microscopy images (projection from z-stacks) showing a phagocytic react-NSC (GFP+, GFAP+) engulfing an apoptotic cell (KA 7d). A high magnification orthogonal view is shown in the middle panel. A side orthogonal view of the phagocytic react-NSC is shown in the right panel. (E) Quantification of apoptotic cells per mm^3^ in the GCL and SGZ of the DG at 7dpKA. (F) Quantification of the % of phagocytic NSCs at 7dpKA.

To test whether NSC phagocytosis was occurring at later time points after KA injection, we assessed NSC phagocytosis at 7dpPBS/KA injection in LPA_1_-GFP mice. As in the 3dpKA time point, we quantified the phagocytosis of apoptotic cells by NSCs and react-NSCs in the GCL and SGZ of the DG (Figure 1D). Our quantifications show a great increase in apoptotic numbers at 7dpKA (Figure 1E) and again a 0,10% of phagocytic NSCs at 7dpKA as compared to none in C mice (Figure 1F). However, the difference in phagocytosis of react-NSCs is not significant due to the numbers being really low and the variability between animals being higher than at 3dpKA. In conclusion, this data show that react-NSCs phagocytose apoptotic cells at both 3 and 7dpKA.

### Hippocampal react-NSCs phagocytose apoptotic cells in MTLE and TBI in Nestin-GFP mice

Because LPA_1_-GFP mice have a very low GFP fluorescence, we hypothesized that this could affect the percentage of phagocytic react-NSCs that we could visualize. Additionally, we also wanted to validate our observations in another transgenic mouse strain. Thus, we decided to quantify phagocytosis in Nestin-GFP mice, where GFP is expressed in NSCs although not specifically (Mignone et al. 2004). In these mice, NSCs and react-NSCs can be easily recognized using immunofluorescence and confocal microscopy, due to their co-expression of GFP and GFAP, radial glia morphology, and lack of S100β (Valcárcel-Martín et al. 2020). Thus, we induced our model of MTLE in Nestin-GFP mice, and waited until 3dpPBS/KA. We then proceeded to quantify the phagocytosis of apoptotic cells by NSCs and react-NSCs (Figure 2A). Our quantifications show an overall increase in apoptotic numbers (Figure 2B) and 8% of phagocytic react-NCSs at 3dpKA as compared to no phagocytosis by NSCs in controls (Figure 2C).

**Figure 2.**
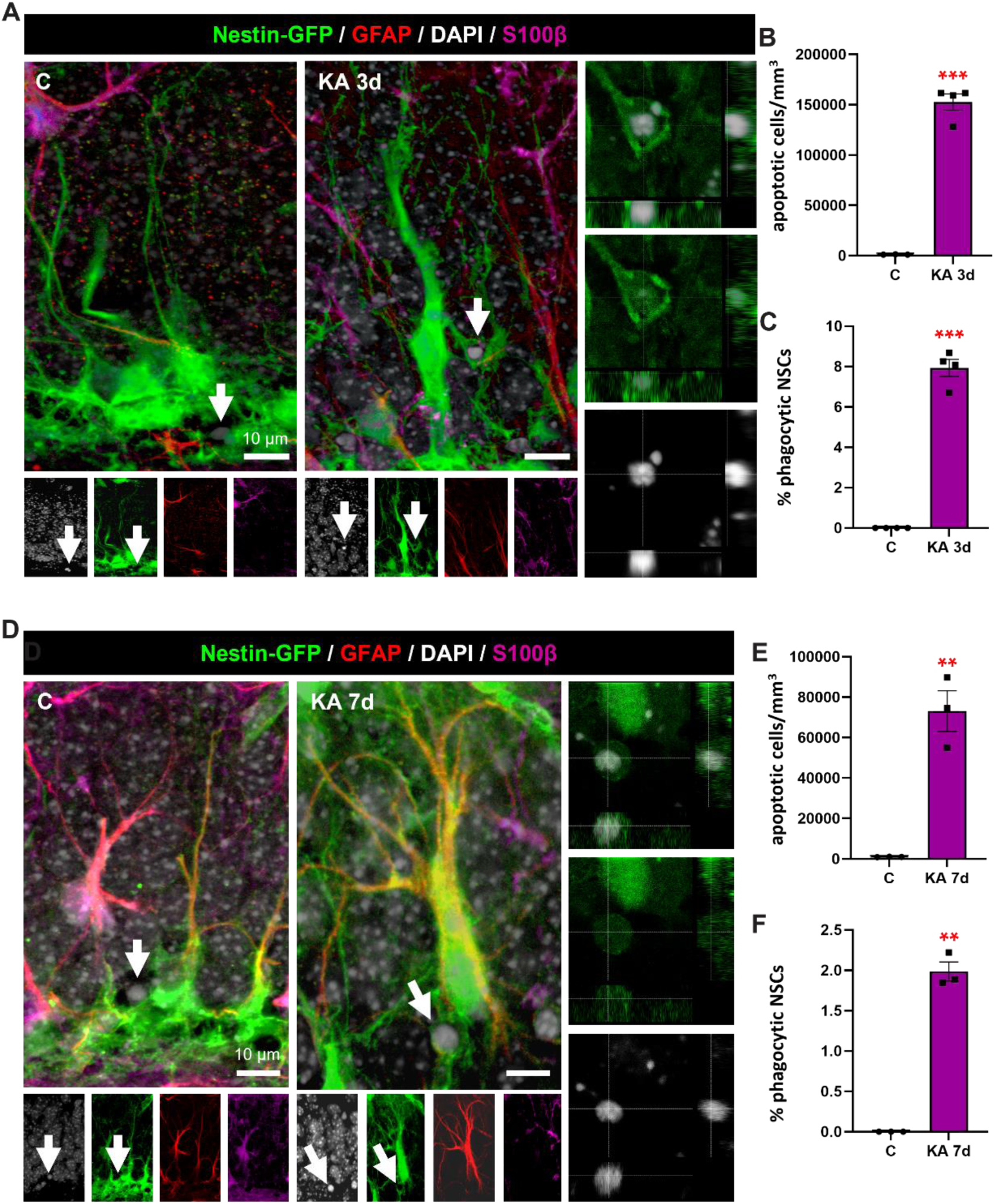
Hippocampal react-NSCs phagocytose apoptotic cells in a model of MTLE in Nestin-GFP mice. (A) Confocal microscopy images (projection from z-stacks) showing a non-phagocytic NSC (GFP+, GFAP+, S100β-) and an apoptotic cell (pyknotic, stained with DAPI) in control and a phagocytic react-NSC (GFP+, GFAP+, S100β-) engulfing an apoptotic cell (KA 3d). A high magnification orthogonal view of the phagocytic pouch is shown in the right panel for KA3d. (B) Quantification of apoptotic cells per mm^3^ in the GCL and SGZ of the DG at 3dpKA. (C) Quantification of the % of phagocytic NSCs at 3dpKA. (D) Confocal microscopy images (projection from z-stacks) showing a non-phagocytic NSC (GFP+, GFAP+, S100β-) and an apoptotic cell (pyknotic, stained with DAPI) in control and a phagocytic react-NSC (GFP+, GFAP+, S100β-) engulfing an apoptotic cell (KA 7d). A high magnification orthogonal view of the phagocytic pouch is shown in the right panel for KA7d. (E) Quantification of apoptotic cells per mm^3^ in the GCL and SGZ of the DG at 7dpKA. (F) Quantification of the % of phagocytic NSCs at 7dpKA.

To test whether NSC phagocytosis was occurring at later time points after KA injection, we then assessed NSC phagocytosis at 7dpPBS/KA injection (Figure 2D). Our quantifications show a great increase in apoptotic cell numbers in 7dpKA (Figure 2E) and a 2% of phagocytic react-NSCs at 7dpKA as compared to none in C mice (Figure 1F). In conclusion, we demonstrate that react-NSCs phagocytose apoptotic cells at both 3 and 7dpPBS/KA in Nestin-GFP mice, proving that this phenomenon is independent of mouse-strain. As predicted, the % of phagocytic react-NSCs are also higher in Nestin-GFP mice, hinting that a higher GFP fluorescence can facilitate the detection of this event.

Additionally, we wanted to test whether react-NSC phagocytosis is specific of MTLE pathology or it could be found in another react-NSC-inducing pathology. For this purpose we resorted to a model of TBI at 14dp impact, where induction of react-NSCs occurs (Figure S1A). Our quantifications show no increase in apoptotic cell numbers in TBI (Figure S1B) and 0,3% of phagocytic react-NCSs at 14dpTBI as compared to no phagocytosis by NSCs in Shams (Figure S1C). In conclusion, these data show that react-NSCs are capable of phagocytosis of apoptotic cells at 14dpTBI and that this event is not tied to a specific pathology. Importantly, react-NSC phagocytosis is not related to the number of apoptotic cells either, as even control condition-like apoptotic cell numbers trigger phagocytosis in react-NSCs but not in NSCs. In conclusion, our data show that react-NSC phagocytosis is an intrinsic capability of these cells.

### Hippocampal react-NSCs show a predominantly immunological RNA signature in early MTLE

We then hypothesized that our observations about increased phagocytosis in react-NSCs in tissue should have a translation in the gene expression of these cells. Thus, we performed a Total RNA sequencing experiment in which we compared NSCs and react-NSCs at 3dp MTLE induction. For this purpose we injected either PBS or KA in the hippocampus of LPA_1_-GFP mice (4 mice per group, 5 samples per treatment) and we specifically isolated the NSC and react-NSC population of the DG based on GFP fluorescence via Fluorescence Activated Cell Sorting (FACS) (Figure 3A). Total RNA sequencing from these cells shows a very high variability in the transcriptomes between control and KA groups and very low intra-group variability as shown by principal component analysis (Figure 3B). Moreover, we performed a Plot MA analysis of Differentially Expressed Genes (DEG) that shows the difference in transcriptome between our two groups (Figure 3C), with 4546 genes upregulated and 3968 genes downregulated in the react-NSC sample.

**Figure 3.**
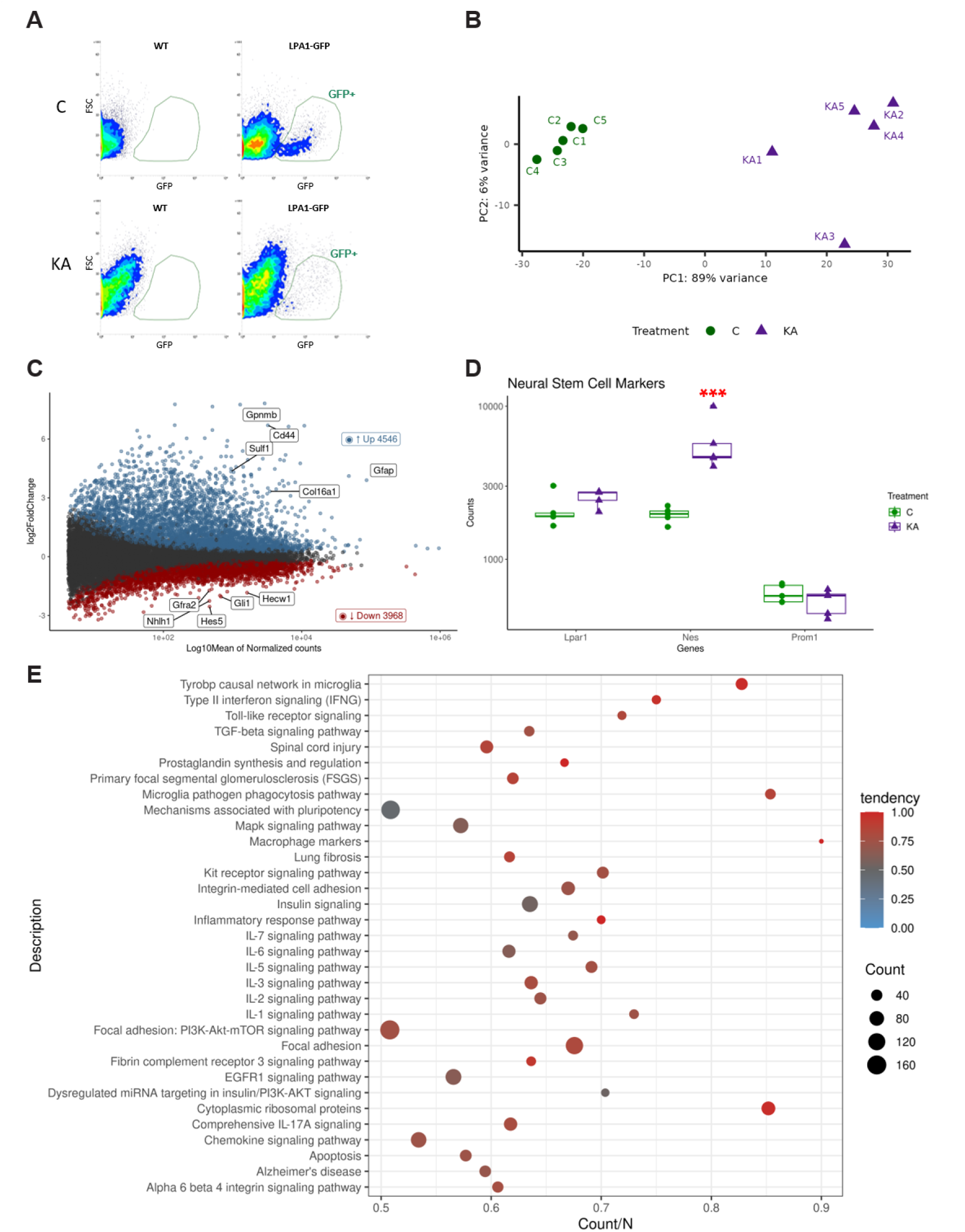
Total RNA sequencing shows a predominant immunological profile of react-NSCs at 3dpKA. (A) Dot plots showing the FACS sorting strategy to isolate NSCs from LPA1-GFP mice based on GFP fluorescence. Dot plots from sorting negative controls (WT mice) are shown on the left and dot plots from LPA1-GFP mice are shown on the right. Only cells from the GFP+ gating (GFP+ NSCs) were sorted for sequencing, after excluding general debris, doublets, and dead/dying cells (data not shown). (B) Principal component analysis (PCA). Reduction dimension technique to summarise transcriptome variability in lower dimensions (X, PCA1; Y, PCA2). Experimental groups are split by the PC1, which summarises 89% of the variance. (C) Plot MA - Differentially expressed genes (DEG). Analysed genes are scattered by its expression levels (axis X, log10mean of Normalised counts) and fold Change (axis Y, log2FC). Differentially expressed genes (padj<0,05) are coloured based on its FC (red, <0; blue >0). (D) Normalised expression of well-known NSC markers. (E) WikiPathways Enrichment Analysis. Enrichment Analysis of all DEG based WikiPathway Ontology. Terms represented are below padj < 0.05, dot size is proportional to the number of DEGs in each GO and the percentage of up and downregulated genes is indicated with color (red and blue, respectively).

To confirm the NSC nature of our two groups we checked the expression of typical NSC markers in the transcriptomic data (Figure 3D). All markers (Nestin, LPAR1 and Prom1) are expressed in both samples, with nestin being significantly more expressed in the KA sample, correlating to the increase we observe in tissue (Sierra et al. 2015). Additionally, we also checked the expression of other progenitor cell markers (Ascl1, Eomes, Ncam1, Neurod1, Neurog2, Pax6) which show lower expression in KA (Figure S2A) and of astrocytic markers (Aldh1I1, GFAP, S100β) which show higher expression in KA (Figure S2B). These data fit with the previously described cellular fate change of react-NSCs, as they get further from their progenitor nature and the neurogenic program and start moving on towards becoming reactive astrocytes (Sierra et al. 2015). Making use of a single cell transcriptomic atlas (Hochgerner et al. 2018), we also predicted the proportion of cell types in our bulk RNA-seq (Figure S2C). This deconvolution confirms the high specificity of the GFP+ NSC isolation protocol, and also hints at a reduction in the percentage of the radial glial cells in favour of adult astrocyte-like cells. These results confirm that the sequenced cells are NSCs and react-NSCs and go in line with the changed fate commitment of react-NSCs we previously described (Sierra et al. 2015).

Finally we wanted to check which were the most differently regulated GOs (gene onthology terms) in react-NSCs. For that purpose we perfomed a WikiPathways enrichment analysis where only the top most differentially expressed GOs are shown (p<0,01) (Figure 3E). We observed that most of the pathways described are immunology-related and are upregulated in react-NSCs, including a pathway involved in “microglia pathogen phagocytosis”. In conclusion, RNA sequencing data confirm the NSC and react-NSC nature of our samples, and show that these cells have very different transcriptomes, with the top differentially expressed genes in react-NSCs showing a predominant immunological profile.

### Hippocampal react-NSCs show a general upregulation of phagocytosis-related genes and pathways in early MTLE

We finally wanted to analyze the RNA sequencing data for differentially regulated phagocytosis-related gene and gene pathway expression. With that purpose we performed an Enriched GO Terms analysis related to phagocytosis (Figure 4A) that shows that several genes in these GOs are differentially expressed in react-NSCs. In order to check whether phagocytosis-related genes and GOs are upregulated in react-NSCs, we performed a Gene Set Enrichment Analysis were all genes relating to “phagocytosis”, “phagocytic vesicles”, and “regulation of phagocytosis” are ordered based on their expression fold change (FC) (Figure 4B). This analysis shows a strong upregulation of the majority of the genes corresponding to these phagocytosis-related pathways in the react-NSC population, which correlates to the phagocytic capacity of react-NSCs that we observe in tissue.

**Figure 4.**
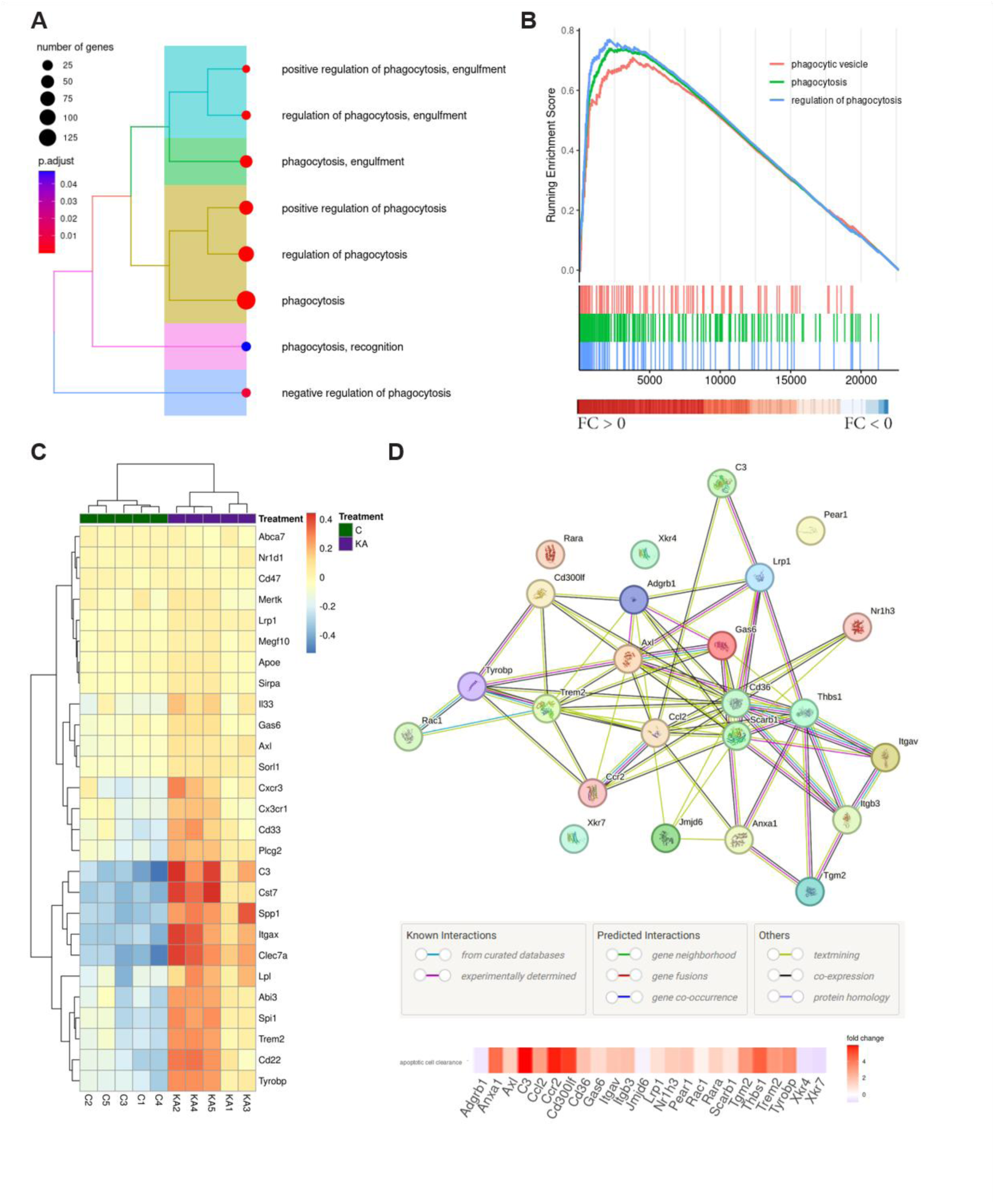
Total RNA sequencing data shows a general upregulation of phagocytosis-related genes and pathways in react-NSCs as 3dpKA. (A) Enriched GO Terms related to phagocytosis. The y axis shows the enriched GOs related to phagocytosis, while the X axis shows how many genes in each category are differentially expressed (Count). The color of each bar matches with its signification (p.adjust). (B) Gene Set Enrichment Analysis of enriched Phagocytic GO Terms. All genes are ordered based on their FC and phagocytosis genes in each GO Terms are highlighted in coloured bars. If a high proportion of bars are located in one end or the other, the running enrichment score will increase. It will be positive if the GO Term is upregulated, as displayed phagocytosis terms, and negative if downregulated. (C) Heatmap Phagocytosis Gene Markers. List of relevant genes related to microglial phagocytosis from literature (Supp Table 1). Genes are clustered based on FC similarities across samples and samples are unbiasedly grouped by its FC across phagocytic genes. (D) Apoptotic Cell Clearance Gene Network. The Network summarises molecular relations among phagocytosis molecules and their respective FC is indicated in the heatmap below.

In order to check for differential expression of specific phagocytosis-related genes, we checked the literature for genes involved in microglial phagocytosis of different kinds of debris (apoptotic cells, myelin, Aβ, etc.) (Table S4) and we performed a heatmap to visualize their up- or down-regulation as shown by their expression fold change (Figure 4C). Results show a clear upregulation in the expression of 59% of the listed genes in react-NSCs. Moreover, we also performed a KEGG Phagosome Pathway analysis of Differentially expressed genes, and a KEGG Toll-like receptor signalling pathway analysis (Figure S3A), to analyse the expression fold change in genes expressed in these pathways. The results show a clear upregulation in many of these pathway genes in react-NSCs.

Finally, we wanted to check whether our specific observation of apoptotic cell phagocytosis was also reflected on gene expression. For that purpose, we performed an apoptotic cell clearance gene network analysis (Figure 4D). The network summarises molecular relations among genes involved in apoptotic cell clearance based on known and predicted interactions and their respective FC is indicated in the heatmap below. This analysis shows that most of the apoptotic cell clearance genes are upregulated in the react-NSC population. In conclusion, these results demonstrate that react-NSCs show differential gene expression and a general upregulation of phagocytosis-related genes, including specific genes related to apoptotic cell clearance.

## DISCUSSION

In this paper we provide evidence of phagocytosis as a new extra-neurogenic function of hippocampal react-NSCs in comparison to physiological NSCs. In a mouse model of MTLE, react-NSCs are capable of phagocytosing apoptotic cells at 3 and 7dpKA in LPA1-GFP mice and in Nestin-GFP mice, showing that this capacity is unrelated to a specific mouse line. Moreover, we also show that react-NSCs that arise in a different pathological model (TBI) are also capable of phagocytosis, showing that this capability is inherent to react-NSCs and not due to a specific pathological model. Importantly, react-NSC phagocytosis is not related to the number of apoptotic cells, as the same number of apoptotic cells trigger phagocytosis in react-NSCs but not in NSCs. Thus, we can conclude that the capability to phagocytose apoptotic cells is a differentiating feature of react-NSCs. Furthermore, sequencing data of react-NSCs at 3dpKA show that these cells have a heightened immunological gene expression profile when compared to NSCs. Importantly, and in accordance with our in-tissue observations, react-NSCs also over-express many phagocytosis-related gene pathways, as well as genes involved in phagosome formation. Thus, our results conclude that react-NSCs have a prominent immunological profile comparing to NSCs, that is justified both in gene expression and functionally by their capability to phagocytose dead cells.

Our group first described react-NSCs in the same MTLE model we have used in this paper. In this model, neuronal hyperactivity causes seizures inducing the change of NSCs into react-NSCs, which become hypertrophic and multibranched, get activated to enter the cell cycle in large numbers, and also switch from asymmetric to symmetric division, giving rise to even more react-NSCs which eventually become reactive astrocytes (Sierra et al. 2015). As a consequence, neurogenesis becomes nearly abolished in the DG. Importantly, react-NSCs have also been found to arise in other pathological models such as TBI. Interestingly, our results regarding stem cell, neural progenitor, and astrocytic markers back up this cell-fate change in react-NSCs that we previously described (Sierra et al. 2015). React-NSCs express less progenitor markers and more astrocytic markers, which goes in line with these cells starting to move towards their final fate of becoming reactive astrocytes.

Our results show that react-NSCs have a prominent immunological profile, where top differentially expressed gene pathways include inflammatory pathways, cytokine signalling, and pathogen phagocytosis. Moreover, most of these genes have an upregulated expression in react-NSCs.

It is well known that the immune system is capable of regulating hippocampal NSCs and adult neurogenesis, both in pathological and in physiological conditions . Among the pathological situations affecting NSCs and neurogenesis inflammation has been the most widely studied, showing that the acute activation of local or systemic innate pro-inflammatory cascades exerts a detrimental impact on postnatal and adult neurogenesis (Carpentier y Palmer 2009).

However, fewer evidence exists for NSCs communicating with their surroundings via immune mediators. It has been reported that NSCs and precursor cells in the brain express a wide range of TLRs, thus sensing a broad range of immunomodulatory signals (Sallustio et al. 2019). Experiments using mice deficient in different TLRs show these receptors have several roles in regulating NSC proliferation, differentiation, and migration (Alvarado y Lathia 2016). These findings align with our results, as we see an upregulation of gene expression in several Toll-like receptors and in their signalling pathway in react-NSCs, which also have different proliferation and differentiation patterns compared to NSCs. Interestingly, chronic inflammation has been found to direct olfactory stem cell functional switch from neuroregeneration to immune defense, shutting down the generation of new sensory neurons and secreting cytokines and chemokines to attract peripheral immune cells (Chen, Reed, y Lane 2019). These results also relate to our findings, as there is also heightened inflammation in the hippocampus in our MTLE model (Abiega et al. 2016) and react-NSCs become astrogliogenic instead of neurogenic and have an overexpression of immune-related genes.

Interestingly, proliferative hippocampal NSCs have previously been linked to immunological activity. The proliferative fraction of NSCs isolated from LPA_1_-GFP mice show an enrichment in immune-response-associated genes when analyzed using bulk RNA-sequencing (Walker et al. 2016). Our data match this data, as react-NSCs are a proliferative population in comparison to NSCs and show a general upregulation of immune-related genes. Another paper reports that Tcf4 deletion causes a decrease in the proliferation of progenitors leading to a reduction of adult neurogenesis in vivo (Shariq et al. 2021). Interestingly, Tcf4 KO in nestin-expressing hippocampal progenitors in vitro decreases their proliferative capacity and makes them acquire myeloid inflammatory characteristics as compared to WT progenitors (Shariq et al. 2021). These conflicting results could be due to the differences between in vivo and in vitro conditions but could also point to the fact that the acquisition of an immunological profile by NSCs might not be exclusive to an upregulation of proliferation but to a dysregulation of this process, be it an increase or a decrease. However, these papers do not report any upregulation of phagocytosis-related genes or GO terms.

In this paper, we describe a new extra-neurogenic function of hippocampal react-NSCs as cells that are capable of phagocytosing apoptotic cells in vivo. There are very few reports on NSCs and phagocytosis, and they mainly involve in vitro experiments and phagocytosis of synthetic phagocytic targets, such as latex beads. Regarding in vitro experiments, it has been previously described that cultured hippocampal NPCs can phagocytose carboxylated beads (Leeson et al. 2018). Both cultured SVZ and hippocampal NPCs show uptake of beads, although it is higher in SVZ NPCs. Another paper also reports that cultured SVZ cells with neural stem-cell features are capable of phagocytosing latex beads and apoptotic cell derived fragments as analyzed by flow cytometry (Ginisty et al. 2015). We report that NSCs don’t perform phagocytosis, which seems to contradict this in vitro data obtained from control NSCs. However, culturing NSCs in vitro can hardly be expected to mimic their physiological in vivo environment and these cells are probably much closer to resembling a pathological condition than a physiological one. There’s only one paper reporting in vivo phagocytosis of SVZ NSCs, where they show that nestin-, Sox2-, and ALDH-expressing neural stem-like cells engulf latex beads or apoptotic cell-derived fragments that were injected into mice lateral brain ventricles (Ginisty et al. 2015). Latex bead phagocytosis was assessed in tissue and apoptotic cell fragment phagocytosis was assessed after tissue dissociation. It is important to note that latex beads represent an indirect measure of phagocytosis, as they don’t release any chemoattractants or express any phagocytic ligands, meaning that they do not activate conventional finding or engulfing mechanisms in phagocytes (Park, Ishihara, y Cox 2011; Diaz-Aparicio et al. 2016). Thus, the lack of resemblance to the in vivo phagocytic process hinders the biological relevance of these experiments (Diaz-Aparicio et al. 2016). Moreover, it is unclear why the authors choose not to assess the phagocytosis of apoptotic cell fragments in tissue. Our results provide the very first direct evidence of hippocampal react-NSCs in vivo phagocytosing a biologically relevant target such as apoptotic cells.

Other cells of the neurogenic niche have been previously described to be able to perform phagocytosis in vivo besides microglia, such as neuroblasts and astrocytes. Astrocytes are capable of phagocytosing a spectrum of neuronal material, including synapses, apoptotic neurons, degenerating axons, amyloid plaques in AD, and α-synuclein in Parkinson’s disease (Lee y Chung 2021). Hippocampal astrocytes can phagocytose synapses in vivo in the hippocampal CA1 (Lee y Chung 2021). Astrocytes are also capable of phagocytosing apoptotic cells in vivo at 1 day post-induction of MTLE via KA (Abiega et al. 2016). Importantly, these results align with our data supporting the cell-fate change of NSCs to become react-NSCs, which express less progenitor markers and more astrocytic markers, showing that these cells are closer to astrocytes and could also be functionally closer to them in terms of their capability for phagocytosis. Whether react-NSCs can phagocytose other types of targets such as synapses and the functional consequences of this process for the tissue still remain to be elucidated.

In conclusion, our data describe a novel extra-neurogenic function of react-NSCs as cells capable of performing a canonical immune function as phagocytosis of apoptotic cells. Our results further portray react-NSCs as cells that are not just passive responders to immune signalling but acquire an immunological gene expression profile, making them potentially involved in immune crosstalk. Comprehending the interaction between react-NSCs and the immune system and the consequences of this will advance our understanding of their biology and potentially contribute to their modulation in pathologies like epilepsy and TBI.

## ACKNOWLEDGEMENTS

We thank all the members of the Laboratory of Neurogenesis, Neuroinflammation and Network Dynamics. OAE was supported by the Basque Government (postdoctoral fellowship and HTC-AI IKUR). RSG holds a predoctoral fellowship from the Tatiana Perez de Guzman el Bueno Foundation. TMG is a recipient of a predoctoral fellowship from the Basque Government. LRC is a recipient of a predoctoral fellowship from the Basque Government. FGM was supported by IKERBASQUE, the Spanish Ministry (PID2021-125156NB-I00, MINECO/MICINN) and the Basque Government (PIBA_2022_1_0027). CPF was supported by Alzheimer Nederland (WE. 03-2020-0). SMS was supported by Ramón y Cajal Program (RyC 2021-033215-I, MINECO/MICINN) and the Basque Government (PIBA_2023_1_0045). JME was supported by the Spanish Ministry of Economy and Competitiveness/Innovation and Science (MINECO/MICINN) Ramón y Cajal Program (RyC 2012-11137; PCIN-2016-128), ERA-NET-NEURON III program, PID2019-104766RB-C21 and CPP2022-009779, Basque Government (PIBA_2021_1_0018), La Caixa Foundation Health Research grant (HR23-00860); Fundación Alicia Koplowitz and Federación Española de Enfermedades Raras (FEDER). We thank the Cell Analytics Facility of Achucarro and specially Dr. Laura Escobar (Leioa, Spain) for her help with the cell sorting experiments. We thank Dr. Ana María Aransay head of the the Platform for Genome Analyses of CIC bioGUNE (Derio, Spain) for her help with the RNA sequencing experiments.

## AUTHOR CONTRIBUTIONS

OA designed and performed experiments, analysed and interpreted results, and wrote the manuscript. RSG analysed and interpreted results. MEO, and SMS performed experiments and analysed results. TMG and LRC performed experiments. FGM interpreted results. CPF designed experiments. J.M.E. conceived the project, designed experiments, interpreted results, and reviewed the manuscript.

