## Supplementary material for "Hippocampal reactive neural stem cells are able to phagocytose and have an immunological molecular signature": Figure S1A: SUPPLEMENTARY FIGURES final.docx

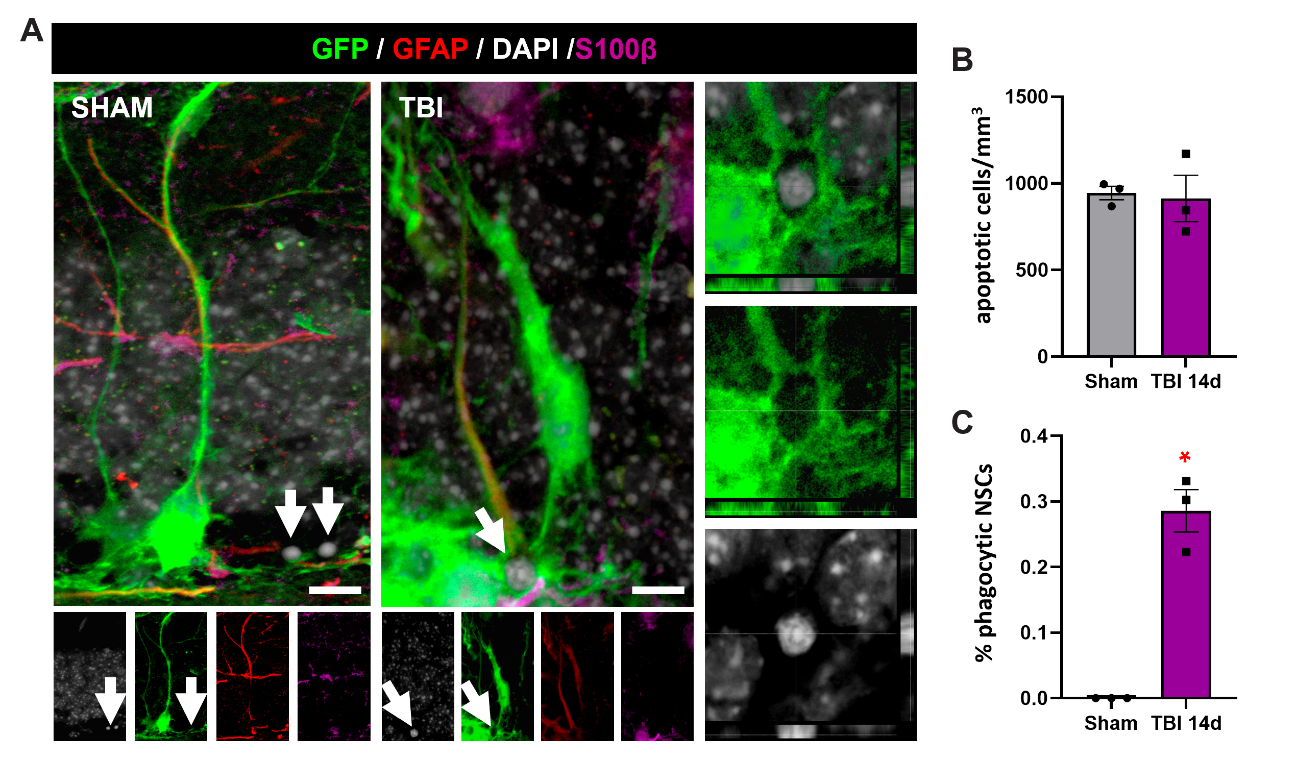

**Figure S1. Hippocampal react-NSCs phagocytose apoptotic cells in a model of TBI in Nestin-GFP mice.** (a) Confocal microscopy images (projection from z‐stacks) showing a non-phagocytic NSC (GFP+, GFAP+) and an apoptotic cell (pyknotic, stained with DAPI) in Sham and a phagocytic react-NSC (GFP+, GFAP+) engulfing an apoptotic cell (TBI 14d). A high magnification orthogonal view of the phagocytic pouch is shown in the right panel for TBI. (b) Quantification of apoptotic cells per mm^3^ in the GCL and SGZ of the DG at 14dpTBI. (c) Quantification of the % of phagocytic NSCs at 14dpTBI.

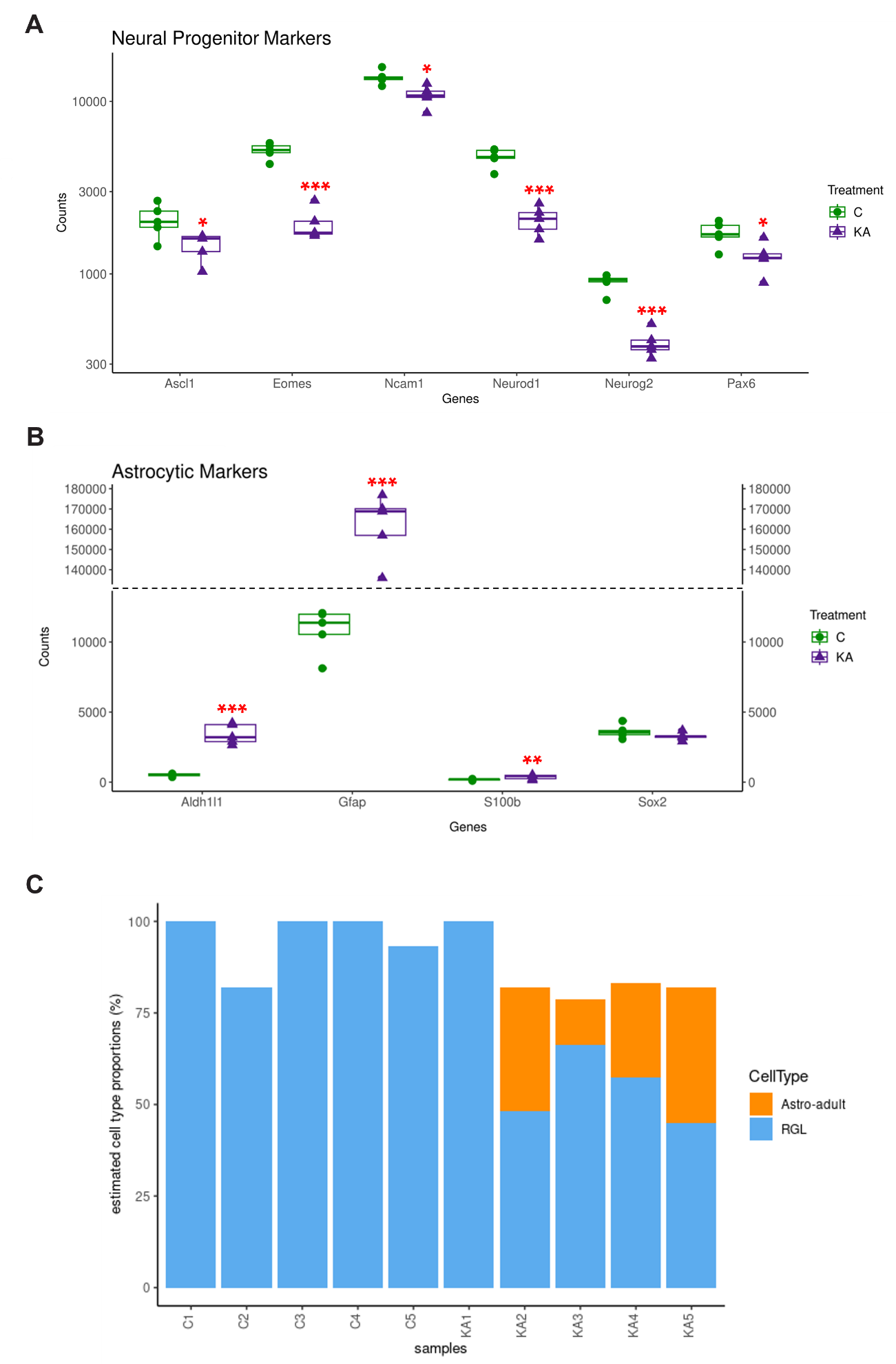

**Figure S2. Differential expression of cell identity genes in react-NSCs.** (a) Normalised expression of well-known progenitor cell markers. (b) Normalised expression of well-known astrocytic markers. (c) Cell Type Estimation in RNA-seq samples. The RNA-seq deconvolution (svr method/granulator package) generates coloured bars indicating the proportion of estimated known cell type: astrocytes (orange) and RGL (radial glia, blue) in each bulk sample based on the single cell transcriptomic dataset of Haring et al. 2018.

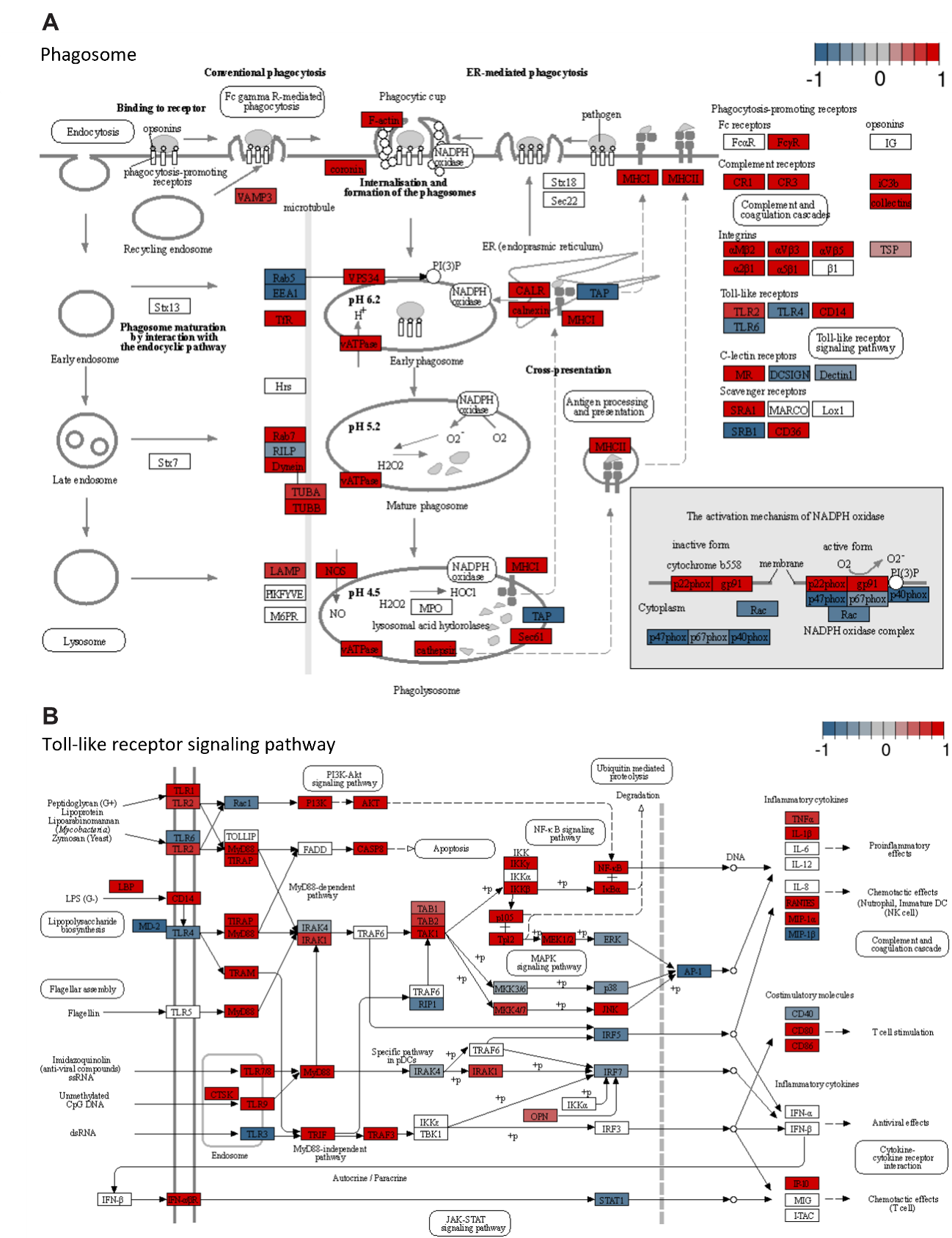

**Figure S3. Genes involded in phagosome formation and in Toll-like receptor signaling show a general upregulation in react-NSCs.** (a) KEGG Phagosome Pathway, differentially expressed genes are coloured, red for positive FC, blue for negative FC. (b) KEGG Toll-like receptor signalling Pathway, differentially expressed genes are coloured, red for positive FC, blue for negative FC.

| **Phagocytosis related gene** | **Phagocytosed debris type** | **Phagocytosis regulation (Pos/Neg)** | **Condition of altered expression** | **References** |
| --- | --- | --- | --- | --- |
| Abca7 | Aβ | Pos | AD | (Fu et al. 2016), (Jonas, Jain, y Li 2022) |
| ABI3 | Aβ | Pos | AD, aging | (Butler et al. 2021; Karahan et al. 2023) |
| Apoe | Apoptotic cells | Pos | AD | (Butler et al. 2021; Muth et al. 2019) |
| Axl | Apoptotic cells | Pos | PD | (Butler et al. 2021; Fourgeaud et al. 2016) |
| C3 | Synapses | Pos | MS | (Li et al. 2022; Michailidou et al. 2015) |
| CD22 | Myelin, Aβ, α-synuclein | Neg | Aging | (Pluvinage et al. 2019) |
| CD33 | Aβ, myelin, dextran | Pos | AD | (Butler et al. 2021; Bhattacherjee et al. 2021) |
| CD47 | Synapses | Neg | AD | (Li et al. 2022; Ding et al. 2021) |
| Clec7α | Beta-glucan exposing cells | Pos | N/A | (Butler et al. 2021) |
| Cst7 | N/A | Neg | AD | (G. S. Kim et al. 2023; Daniels et al., s. f.) |
| CX3CR1 | Synapses | Pos | Whisker lesion | (Li et al. 2022; Gunner et al. 2019) |
| CXCR3 | N/A | Neg | LPS induced inflammation | (Li et al. 2022; De Jong et al. 2008) |
| IL-33/ST2 | Extracellular matrix, Aβ, hematoma | Pos | N/A | (Li et al. 2022; Nguyen et al. 2020; He et al. 2022; Lau et al. 2020; Xu et al. 2020) |
| Itgax | iC3b-opsonized cells | Pos | DAM | (Butler et al. 2021) |
| LpI | Aβ | Pos | DAM, AD | (G. S. Kim et al. 2023; Loving y Bruce 2020) |
| LRP1 | Aβ | Pos | AD | (Butler et al. 2021; Loving y Bruce 2020) |
| MEGF10 | Neurons, apoptotic cells, synapses | Pos | Stroke | (Butler et al. 2021) |
| MERTK | Neurons | Pos | Cerebral ischemia | (Butler et al. 2021; Neher et al. 2013) |
| PLCG2 | Aβ | Pos/Neg depending on variant | AD | (Butler et al. 2021; Tsai et al. 2023) |
| SIRPα | Synapses, neurons, myelin | Neg | AD | (Butler et al. 2021; Ding et al. 2021; Ransohoff y Perry 2009; Noda y Suzumura 2012) |
| SORL1 | Aβ | Pos | AD | (Podleśny-Drabiniok, Marcora, y Goate 2020; Mishra et al. 2022) |
| SPI1/PU.1 | Aβ | Pos | AD | (Podleśny-Drabiniok, Marcora, y Goate 2020; Smith et al. 2013) |
| Spp1 | Aβ | Pos | AD | (G. S. Kim et al. 2023; De Schepper et al. 2023) |
| TREM2 | Synapses, neurons, Aβ, myelin, PS exposing cells | Pos | AD, MS | (Li et al. 2022; Butler et al. 2021; S.-M. Kim et al. 2017; Atagi et al. 2015; Takahashi, Rochford, y Neumann 2005; Leyns et al. 2017; Filipello et al. 2018; Cignarella et al. 2020) |
| Tyrobp | Aβ | Pos | AD | (G. S. Kim et al. 2023; Haure-Mirande et al. 2017) |

Table 1: Genes related to microglial phagocytosis. Abbreviations: Pos, positive; Neg, negative; N/A, not available; Aβ, amyloid β; PS, phosphatidylserine; AD, Alzheimer’s disease; PD, Parkinson’s disease; MS, multiple sclerosis; LPS, lipopolysaccharide; DAM, disease-associated microglia; ABCA7, ATP-binding cassette sub-family A member 7; ABI3; ABI family member 3; ApoE, apolipoprotein E; C3, complement C3; Clec7a, C-Type Lectin Domain Containing 7A; Cst7, Cystatin F; CX3CR1, C-X3-C Motif Chemokine Receptor 1; IL-33/ST2, Interleukin 33; Itgax, Integrin alpha-X; Lpl, lipoprotein lipase; LRP1, LDL Receptor Related Protein 1; MEGF10, multiple EGF like domains 10; MERTK, Mer receptor tyrosine kinase, Tyrosine Kinase; PLCG2, phospholipase C gamma 2; SIRPα, Signal Regulatory Protein Alpha; SORL1, sortilin related receptor 1; SPI1/PU.1, transcription factor PU.1; Spp1, Secreted Phosphoprotein 1; TREM2, triggering receptor expressed on myeloid cells 2; Tyrobp, TYRO protein tyrosine kinase binding protein.

Daniels, Michael JD, Lucas Lefevre, Stefan Szymkowiak, Alice Drake, Laura McCulloch, Makis Tzioras, Jack Barrington, et al. s. f. «Cystatin F (Cst7) drives sex-dependent changes in microglia in an amyloid-driven model of Alzheimer’s disease». *eLife* 12: e85279. https://doi.org/10.7554/eLife.85279.

De Jong, Eiko K., Alexander H. De Haas, Nieske Brouwer, Hilmar R. J. Van Weering, Marjolein Hensens, Ingo Bechmann, Pierre Pratley, Evelyn Wesseling, Hendrikus W. G. M. Boddeke, y Knut Biber. 2008. «Expression of CXCL4 in Microglia in Vitro and in Vivo and Its Possible Signaling through CXCR3». *Journal of Neurochemistry* 105 (5): 1726-36. https://doi.org/10.1111/j.1471-4159.2008.05267.x.

De Schepper, Sebastiaan, Judy Z. Ge, Gerard Crowley, Laís S. S. Ferreira, Dylan Garceau, Christina E. Toomey, Dimitra Sokolova, et al. 2023. «Perivascular Cells Induce Microglial Phagocytic States and Synaptic Engulfment via SPP1 in Mouse Models of Alzheimer’s Disease». *Nature Neuroscience* 26 (3): 406-15. https://doi.org/10.1038/s41593-023-01257-z.

Ding, Xin, Jin Wang, Miaoxin Huang, Zhangpeng Chen, Jing Liu, Qipeng Zhang, Chenyu Zhang, Yang Xiang, Ke Zen, y Liang Li. 2021. «Loss of Microglial SIRPα Promotes Synaptic Pruning in Preclinical Models of Neurodegeneration». *Nature Communications* 12 (1): 2030. https://doi.org/10.1038/s41467-021-22301-1.

Filipello, Fabia, Raffaella Morini, Irene Corradini, Valerio Zerbi, Alice Canzi, Bernadeta Michalski, Marco Erreni, et al. 2018. «The Microglial Innate Immune Receptor TREM2 Is Required for Synapse Elimination and Normal Brain Connectivity». *Immunity* 48 (5): 979-991.e8. https://doi.org/10.1016/j.immuni.2018.04.016.

Fourgeaud, Lawrence, Paqui G. Través, Yusuf Tufail, Humberto Leal-Bailey, Erin D. Lew, Patrick G. Burrola, Perri Callaway, et al. 2016. «TAM Receptors Regulate Multiple Features of Microglial Physiology». *Nature* 532 (7598): 240-44. https://doi.org/10.1038/nature17630.

Fu, YuHong, Jen-Hsiang T. Hsiao, George Paxinos, Glenda M. Halliday, y Woojin Scott Kim. 2016. «ABCA7 Mediates Phagocytic Clearance of Amyloid-β in the Brain». *Journal of Alzheimer’s Disease: JAD* 54 (2): 569-84. https://doi.org/10.3233/JAD-160456.

Gunner, Georgia, Lucas Cheadle, Kasey M. Johnson, Pinar Ayata, Ana Badimon, Erica Mondo, M. Aurel Nagy, et al. 2019. «Sensory Lesioning Induces Microglial Synapse Elimination via ADAM10 and Fractalkine Signaling». *Nature Neuroscience* 22 (7): 1075-88. https://doi.org/10.1038/s41593-019-0419-y.

Haure-Mirande, Jean-Vianney, Mickael Audrain, Tomas Fanutza, Soong Ho Kim, William L. Klein, Charles Glabe, Ben Readhead, et al. 2017. «Deficiency of TYROBP, an Adapter Protein for TREM2 and CR3 Receptors, Is Neuroprotective in a Mouse Model of Early Alzheimer’s Pathology». *Acta Neuropathologica* 134 (5): 769-88. https://doi.org/10.1007/s00401-017-1737-3.

He, Danyang, Heping Xu, Huiyuan Zhang, Ruihan Tang, Yangning Lan, Ruxiao Xing, Shaomin Li, et al. 2022. «Disruption of the IL-33-ST2-AKT Signaling Axis Impairs Neurodevelopment by Inhibiting Microglial Metabolic Adaptation and Phagocytic Function». *Immunity* 55 (1): 159-173.e9. https://doi.org/10.1016/j.immuni.2021.12.001.

Jonas, Lauren A., Tanya Jain, y Yue-Ming Li. 2022. «Functional insight into LOAD-associated microglial response genes». *Open Biology* 12 (1): 210280. https://doi.org/10.1098/rsob.210280.

Karahan, Hande, Daniel C. Smith, Byungwook Kim, Brianne McCord, Jordan Mantor, Sutha K. John, Md Mamun Al-Amin, Luke C. Dabin, y Jungsu Kim. 2023. «The effect of Abi3 locus deletion on the progression of Alzheimer’s disease-related pathologies». *Frontiers in Immunology* 14. https://www.frontiersin.org/articles/10.3389/fimmu.2023.1102530.

Kim, Gab Seok, Elisabeth Harmon, Manuel Gutierrez, Jessica Stephenson, Anjali Chauhan, Anik Banerjee, Zachary Wise, et al. 2023. «Single-cell analysis identifies Ifi27l2a as a novel gene regulator of microglial inflammation in the context of aging and stroke». *Research Square*, febrero, rs.3.rs-2557290. https://doi.org/10.21203/rs.3.rs-2557290/v1.

Kim, Su-Man, Bo-Ram Mun, Sun-Jun Lee, Yechan Joh, Hwa-Youn Lee, Kon-Young Ji, Ha-Rim Choi, et al. 2017. «TREM2 Promotes Aβ Phagocytosis by Upregulating C/EBPα-Dependent CD36 Expression in Microglia». *Scientific Reports* 7 (1): 11118. https://doi.org/10.1038/s41598-017-11634-x.

Lau, Shun-Fat, Congping Chen, Wing-Yu Fu, Jianan Y. Qu, Tom H. Cheung, Amy K. Y. Fu, y Nancy Y. Ip. 2020. «IL-33-PU.1 Transcriptome Reprogramming Drives Functional State Transition and Clearance Activity of Microglia in Alzheimer’s Disease». *Cell Reports* 31 (3). https://doi.org/10.1016/j.celrep.2020.107530.

Leyns, Cheryl E. G., Jason D. Ulrich, Mary B. Finn, Floy R. Stewart, Lauren J. Koscal, Javier Remolina Serrano, Grace O. Robinson, Elise Anderson, Marco Colonna, y David M. Holtzman. 2017. «TREM2 Deficiency Attenuates Neuroinflammation and Protects against Neurodegeneration in a Mouse Model of Tauopathy». *Proceedings of the National Academy of Sciences of the United States of America* 114 (43): 11524-29. https://doi.org/10.1073/pnas.1710311114.

Li, Congqin, Yong Wang, Ying Xing, Jing Han, Yuqian Zhang, Anjing Zhang, Jian Hu, Yan Hua, y Yulong Bai. 2022. «Regulation of microglia phagocytosis and potential involvement of exercise». *Frontiers in Cellular Neuroscience* 16. https://www.frontiersin.org/articles/10.3389/fncel.2022.953534.

Loving, Bailey A., y Kimberley D. Bruce. 2020. «Lipid and Lipoprotein Metabolism in Microglia». *Frontiers in Physiology* 11. https://www.frontiersin.org/articles/10.3389/fphys.2020.00393.

Michailidou, Iliana, Janske G. P. Willems, Evert-Jan Kooi, Corbert van Eden, Stefan M. Gold, Jeroen J. G. Geurts, Frank Baas, Inge Huitinga, y Valeria Ramaglia. 2015. «Complement C1q-C3–Associated Synaptic Changes in Multiple Sclerosis Hippocampus». *Annals of Neurology* 77 (6): 1007-26. https://doi.org/10.1002/ana.24398.

Mishra, Swati, Allison Knupp, Jessica E. Young, y Suman Jayadev. 2022. «Depletion of the AD Risk Gene SORL1 Causes Endo-Lysosomal Dysfunction in Human Microglia». *Alzheimer’s & Dementia* 18 (S4): e068943. https://doi.org/10.1002/alz.068943.

Muth, Christiane, Alexander Hartmann, Diego Sepulveda-Falla, Markus Glatzel, y Susanne Krasemann. 2019. «Phagocytosis of Apoptotic Cells Is Specifically Upregulated in ApoE4 Expressing Microglia in vitro». *Frontiers in Cellular Neuroscience* 13. https://www.frontiersin.org/articles/10.3389/fncel.2019.00181.

Neher, Jonas J., Julius V. Emmrich, Michael Fricker, Palwinder K. Mander, Clotilde Théry, y Guy C. Brown. 2013. «Phagocytosis executes delayed neuronal death after focal brain ischemia». *Proceedings of the National Academy of Sciences* 110 (43): E4098-4107. https://doi.org/10.1073/pnas.1308679110.

Nguyen, Phi T., Leah C. Dorman, Simon Pan, Ilia D. Vainchtein, Rafael T. Han, Hiromi Nakao-Inoue, Sunrae E. Taloma, et al. 2020. «Microglial Remodeling of the Extracellular Matrix Promotes Synapse Plasticity». *Cell* 182 (2): 388-403.e15. https://doi.org/10.1016/j.cell.2020.05.050.

Noda, Mariko, y Akio Suzumura. 2012. «Sweepers in the CNS: Microglial Migration and Phagocytosis in the Alzheimer Disease Pathogenesis». *International Journal of Alzheimer’s Disease* 2012 (mayo): e891087. https://doi.org/10.1155/2012/891087.

Pluvinage, John V., Michael S. Haney, Benjamin A. H. Smith, Jerry Sun, Tal Iram, Liana Bonanno, Lulin Li, et al. 2019. «CD22 Blockade Restores Homeostatic Microglial Phagocytosis in Ageing Brains». *Nature* 568 (7751): 187-92. https://doi.org/10.1038/s41586-019-1088-4.

Podleśny-Drabiniok, Anna, Edoardo Marcora, y Alison M. Goate. 2020. «Microglial Phagocytosis: A Disease-Associated Process Emerging from Alzheimer’s Disease Genetics». *Trends in Neurosciences* 43 (12): 965-79. https://doi.org/10.1016/j.tins.2020.10.002.

Ransohoff, Richard M., y V. Hugh Perry. 2009. «Microglial Physiology: Unique Stimuli, Specialized Responses». *Annual Review of Immunology* 27 (1): 119-45. https://doi.org/10.1146/annurev.immunol.021908.132528.

Smith, Amy M., Hannah M. Gibbons, Robyn L. Oldfield, Peter M. Bergin, Edward W. Mee, Richard L. M. Faull, y Mike Dragunow. 2013. «The Transcription Factor PU.1 Is Critical for Viability and Function of Human Brain Microglia». *Glia* 61 (6): 929-42. https://doi.org/10.1002/glia.22486.

Takahashi, Kazuya, Christian D. P. Rochford, y Harald Neumann. 2005. «Clearance of Apoptotic Neurons without Inflammation by Microglial Triggering Receptor Expressed on Myeloid Cells-2». *The Journal of Experimental Medicine* 201 (4): 647-57. https://doi.org/10.1084/jem.20041611.

Tsai, Andy P., Chuanpeng Dong, Peter Bor-Chian Lin, Adrian L. Oblak, Gonzalo Viana Di Prisco, Nian Wang, Nicole Hajicek, et al. 2023. «Genetic Variants of Phospholipase C-Γ2 Alter the Phenotype and Function of Microglia and Confer Differential Risk for Alzheimer’s Disease». *Immunity* 56 (9): 2121-2136.e6. https://doi.org/10.1016/j.immuni.2023.08.008.

Xu, Jing, Zhouqing Chen, Fang Yu, Huan Liu, Cheng Ma, Di Xie, Xiaoming Hu, et al. 2020. «IL-4/STAT6 signaling facilitates innate hematoma resolution and neurological recovery after hemorrhagic stroke in mice». *Proceedings of the National Academy of Sciences* 117 (51): 32679-90. https://doi.org/10.1073/pnas.2018497117.
